# Functionally distinct T-helper cell phenotypes predict resistance to different types of parasites in a wild mammal

**DOI:** 10.1101/2021.03.15.435455

**Authors:** Yolanda Corripio-Miyar, Adam Hayward, Hannah Lemon, Amy R. Sweeny, Xavier Bal, Fiona Kenyon, Jill G Pilkington, Josephine M. Pemberton, Daniel H Nussey, Tom N McNeilly

## Abstract

1. The adaptive immune system is critical to an effective, long-lasting ability to respond to infection in vertebrates and T-helper (Th) cells play a key role in orchestrating the adaptive immune response. Laboratory studies show that functionally distinct Th responses provide protection against different kinds of parasites (i.e., Th1 responses against microparasites and Th2 against macroparasites).
2. Natural populations must deal with challenges from a wide range of infectious agents and co-infection with different types of parasite is the norm, so different Th responses are likely to play an important and dynamic role in maintaining host health and fitness. However, the relationship between T helper immune phenotypes and infection with different types of parasites remains poorly understood in wild animals.
3. In this study, we characterised variation in functionally distinct Th responses (Th1, Th2, Th17 and regulatory responses) in a wild population of Soay sheep using flow cytometry to detect Th-subset specific transcription factors, and *ex vivo* lymphocyte stimulation to quantify release of Th-associated cytokines. We specifically tested the prediction that raised Th1 and Th2 responses should predict reduced apicomplexan (coccidian) and helminth (nematode) parasite burdens, respectively.
4. Cell counts of different Th subsets measured by flow cytometry did not vary with age or sex. However, all measures of Th-associated *ex vivo* cytokine production increased with age, and Th17- and regulatory Th-associated cytokine production increased more rapidly with age in males than females.
5. Independent of age and sex, Th2-associated immune measures negatively predicted gastro-intestinal strongyle nematode faecal egg count, while production of the Th1-associated cytokine IFN-γ negatively predicted coccidian faecal oocyst count.
6. Our results provide important support from outside the laboratory that Th1 and Th2 responses confer resistance to different kinds of parasites (micro- and macro-parasites, respectively). They also add to mounting evidence from wild populations that Th1/Th2 trade-offs often observed in controlled laboratory experiments may not readily translate to more complex natural systems.
7. Our study illustrates that harnessing more specific reagents and tools from laboratory immunology has the potential to illuminate our understanding of epidemiology and host-parasite co-evolution in the wild.

## INTRODUCTION

Research in cellular, medical and veterinary immunology shows us that the vertebrate immune system is highly complex and composed of many different cell types with different functional roles in any particular response to infection (Cox 2001; Coughlan & Lambe 2015; McRae *et al.* 2015; Abolins *et al.* 2017). Variation in the relative abundance of different immune cell types and their responsiveness to stimulation is thought to have major implications for infection risk and disease outcomes (Segerstrom & Miller 2004; Seder, Darrah & Roederer 2008; Albert-Vega *et al.* 2018). The immune system is thought to play an important role in the evolutionary and ecological dynamics of natural vertebrate populations, protecting individuals from infection by a diverse array of parasites and pathogens. Despite this, a shortcoming of many field studies of immunity to date has been the inability to quantify variation in these functionally different cell types, relying often instead on general and generic measures of immune response, such as skin swelling or natural antibody responses to a challenge with a novel antigen (Demas *et al.* 2011; Pedersen & Babayan 2011). In large part this has been due to a lack of a suitable immunological tool kit in non-model systems that would allow the abundance and functionality of important classes of immune cells to be quantified. Recent studies applying tools developed for laboratory rodents to their wild counterparts illustrate striking differences between the immune phenotypes of wild and captive animals (Abolins *et al.* 2011; Abolins *et al.* 2017), as well as variation in diverse aspects of immunity among and within natural populations (Abolins *et al.* 2018). At the same time, studies in wild rodents, rabbits and ungulates highlight the potential for functional constraints on immune-mediated resistance to different types of parasites to influence patterns of infection and disease dynamics in nature (Ezenwa 2016). In order to determine how natural selection has shaped immune responses in wild populations, we first need to characterise variation in functionally-relevant aspects of the immune response, in a way that begins to reflect some of its complexity. Here, we utilise expertise and reagents from veterinary immunology to examine variation in functionally distinct arms of the adaptive immune response and test how this variation predicts abundance of gastrointestinal parasites in wild Soay sheep *(Ovis aries).*

The immune system in vertebrates broadly consists of the innate and adaptive arms (Murphy *et al.* 2012). Innate immune responses are generally non-specific and are rapidly activated immediately after parasite antigens are encountered. In contrast, adaptive immune responses are slower to develop but are highly antigen-specific due to the recognition of specific epitopes within antigens by surface-expressed receptors on B and T lymphocytes. Following antigen-specific activation, B and T cells undergo clonal expansion to generate effector cells that control the immediate infection, and memory cells that provide long-lasting immune protection (Parkin & Cohen 2001; Murphy *et al.* 2012). The adaptive immune system is further divided based on effector functions into humoral immunity, mediated by antibodies produced by B cells, and cellular immunity such as that mediated by cytotoxic T cells and phagocytes (Parkin & Cohen 2001; Murphy *et al.* 2012). Functional diversity of the adaptive immune system is coordinated by CD4^+^ T helper (Th) cells, which can elicit functionally-distinct types of response which support protection of the host against different kinds of challenges (Mosmann & Coffman 1989; Nakayamada *et al.* 2012).

The different types and functions of T helper-mediated immune responses are illustrated in Figure 1. T helper type 1 (Th1) cells promote cellular immune responses to control intracellular pathogens such as viruses and bacteria (Mosmann & Coffman 1989; O’Garra & Robinson 2004). T helper type 2 (Th2) cells promote humoral immunity and are important for controlling large extracellular pathogens including parasitic helminths (Mosmann & Coffman 1989; Romagnani 2000; Grencis 2015). The more recently discovered T helper type (Th17) cells play a role in controlling extracellular bacterial and fungal infections, and are particularly important at mucosal barriers (Stockinger & Omenetti 2017; Sandquist & Kolls 2018). In addition to these effector subsets, CD4^+^ T cells can also differentiate into regulatory T (Treg) cells which play a key role in immune homeostasis and prevention of immunopathology (Pereira *et al.* 2017). Th polarisation is largely decided during the early stages of activation of naïve CD4^+^ T cells, when dendritic cells (specialised antigen presenting cells) and other innate cells ‘sense’ specific pathogen molecules and, through cytokine and other signalling events, trigger unique ‘master’ transcription factors that become lineage-specific markers for the different Th subsets (Figure 1; (Schmitt & Ueno 2015; Gerbe *et al.* 2016; Jain & Pasare 2017)). Each Th subset then secretes a specific group of cytokines that promotes different types of adaptive immune responses, whilst also inhibiting the development of other Th subsets (Seder & Paul 1994; Abbas, Murphy & Sher 1996). Thus, a broad understanding of adaptive immune response function can obtained by quantifying the master transcription factors expressed by Th cells and the cytokines they release upon activation (Figure 1).

**Figure 1.**
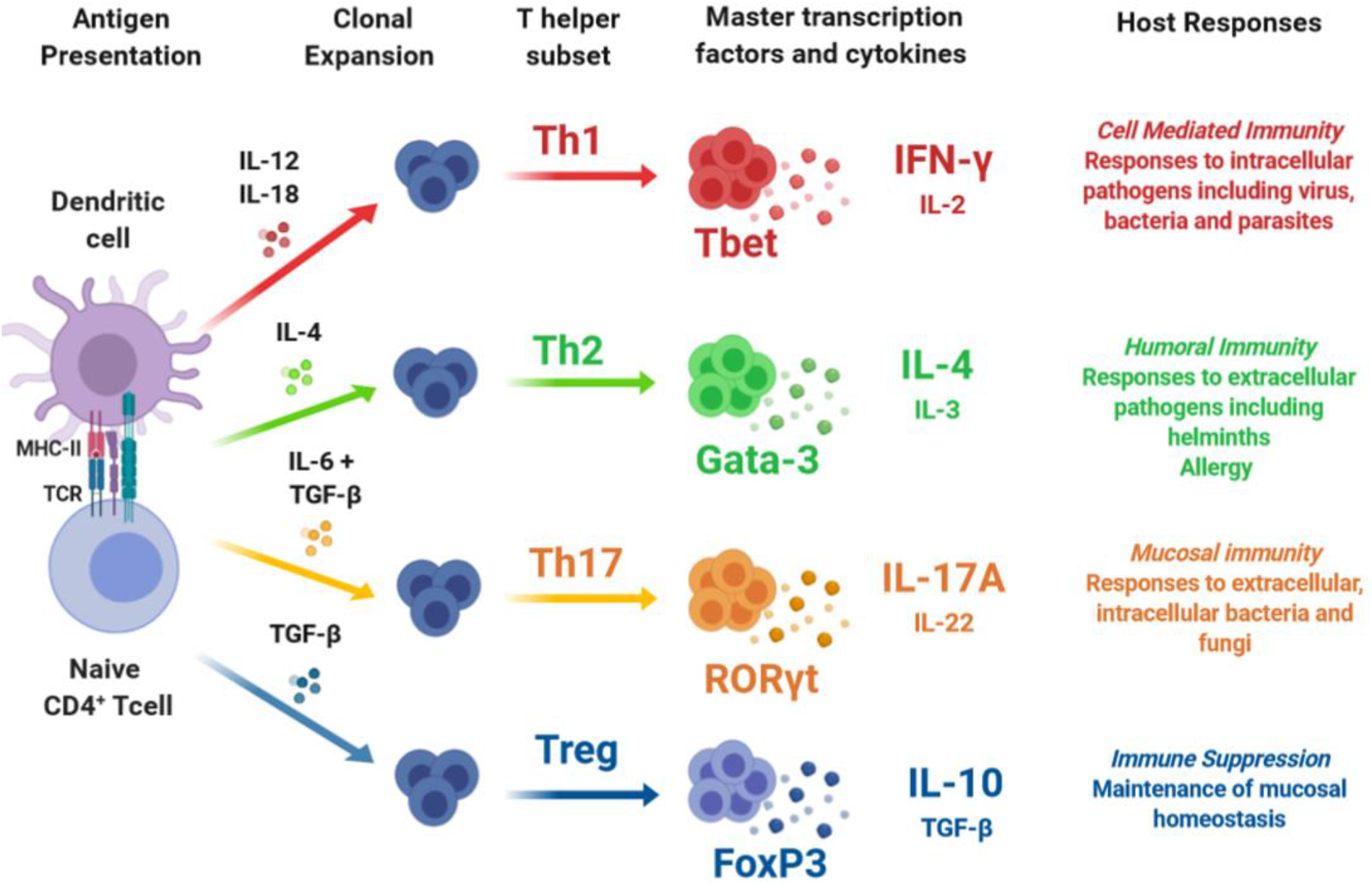
Overview of T helper subsets. Upon encountering foreign antigens, dendritic cells (highly specialised antigen presenting cells) process and present fragments of antigen to naïve CD4^+^ T cells via major histocompatibility complex (MHC) class II molecules. During this process, specific cytokines drive differentiation and clonal expansion of CD4^+^ T cells into functionally distinct T helper (Th) subsets. Each Th subset is associated with a master transcription factor, and secretes specific cytokines involved in coordinating different types of host immune response.

In the wild, vertebrates must deal with challenges from a wide range of infectious agents, and co-infection with many parasites is the norm (Wilson, Fenton & Tompkins 2019). As such, there is good reason to expect that variation in the T helper responses plays a very important role in maintaining host health and fitness under natural conditions. In particular, co-infection with intracellular microparasites (e.g. viruses, bacteria, apicomplexans) and macroparasites (e.g. helminths) are expected to elicit trade-offs between host investment and commitment to Th1 vs. Th2 immune responses (Cox 2001). For example, in experimental helminth infection studies, the greater the suppressive effect of worm infection on Th1 cytokines, the greater the associated increase in microparasite density (Graham 2008). Furthermore, recent studies of wild buffalo show that experimental removal of helminths promotes a Th1 response, with downstream consequences for host resistance to intracellular parasites (Ezenwa *et al.* 2010; Ezenwa & Jolles 2015). However, measurement of functional T cell responses in wild vertebrates is challenging and has rarely been undertaken. As Figure 1 illustrates, measuring an individual’s Th phenotype requires quantification of the number of T cells of different functional types and their functional cytokine response to stimulation. This demands both the immunological tools for T cell phenotyping, which are lacking for most non-model systems, and protocols for either immediate deployment of assays on live cells in the field or careful preservation of cells in the field for later assays (Demas *et al.* 2011). Despite these difficulties, a number of recent studies have begun to explore variation in T cell immunity in wild mammals, by measuring expression of Th master regulator genes in blood samples (Jackson *et al.* 2014; Arriero *et al.* 2017) or by monitoring production of Th1-associated cytokines following *ex vivo* stimulation of lymphocytes derived from blood samples (Beirne, Delahay & Young 2015; Ezenwa & Jolles 2015; Young *et al.* 2020). However, to date, no study in the wild has quantified patterns of variation in the full range of Th phenotypes illustrated in Figure 1 or tested how they predict levels of natural infection with different parasites.

Here, we characterise Th responses in wild Soay sheep (*Ovis aries*) on St Kilda, in which we have previously identified variation in helper T cell and cytotoxic T cell proportions in relation to age and sex (Nussey *et al.* 2012; Watson *et al.* 2016). We use immunological reagents developed for use in domestic sheep to specifically quantify different Th responses (Th1, Th2, Th17 and Treg) through activation-specific cytokine release and expression of Th-associated transcription factors. The Soay sheep on St Kilda are principally infected with two groups of gastrointestinal parasites, nematode worms and coccidian apicomplexans (Craig *et al.* 2006; Craig *et al.* 2008). It is generally accepted that resistance to these parasites groups is broadly associated with different Th responses: Th2 for worms, and Th1 for coccidian parasites (Ovington, Alleva & Kerr 1995; Finkelman *et al.* 2004; Maizels, Hewitson & Smith 2012; Ozmen, Adanir & Haligur 2012; Kim, Chaudhari & Lillehoj 2019), and that, in laboratory settings at least, Th1 and Th2 responses are antagonistic (Seder & Paul 1994; Abbas, Murphy & Sher 1996). We examine the associations among different measures of the four main Th response types and their relationship with age and sex for the first time in the wild. We go on to test the predictions that: (1) worm burdens should be reduced in animals with stronger Th2 responses, (2) coccidia burdens should be reduced in animals with stronger Th1 responses, and (3) Th1 and Th2 responsiveness should trade-off and be negatively correlated.

## MATERIALS AND METHODS

### Study Population

Soay sheep are an ancient breed of domestic sheep that have lived under unmanaged conditions for the last few thousand years in the St Kilda archipelago, 65km west of the Outer Hebrides, Scotland (57°49’N, 08°34’W). In 1932, following the evacuation of the human population from Hirta, the largest island in the archipelago, 107 sheep were moved from the smaller island of Soay onto Hirta (Pemberton & Clutton-Brock 2004). Without human intervention, the population on Hirta has grown to cover the whole island, with numbers only limited by the availability of resources in the grazing grounds. Since 1985, sheep resident to the Village Bay area of Hirta (around a third of the sheep on the island) have been the subject of an intensive individual-based study. Every year, >95% of lambs born in the study area are captured within a week of birth (March-May), given an individual identifying ear tag, weighed and blood- and tissue-sampled for genetic analysis. Each August, as many sheep as possible (~50% on average) are captured using temporary traps in order to collect data on numerous variables including weight and morphometrics, and faecal and blood samples are also collected following Home Office Guidelines (under project license number PPL 70/8818). Samples used in this study were collected from 238 animals captured in August 2019. Blood was collected from sheep by jugular venepuncture into lithium heparin vacutainers (Greiner Bio-One International GmbH) and stored at 4°C prior to being processed. Rectally collected faecal samples were used to estimate strongyle faecal egg counts (FEC) and coccidian faecal oocyst count (FOC) using a modified salt-flotation method (Jackson 1974) as detailed below.

### Whole blood stimulation assays

Within 24h of collection, to examine the cytokine secretion induced by a mitogen by leuocytes, whole blood stimulations were carried out using samples from 208 animals by mixing 1ml of whole blood with 1ml of tissue culture media [RPMI-1640 supplemented with 10% FBS and 50 μM 2-mercaptoethanol, 2 mM L-glutamine, 100 U/ml Penicillin and 100 μg/ml Streptomycin, 5μg/ml Gentamicin (all from Sigma-Aldrich, UK)] containing 10μg/ml final concentration of poke weed mitogen (PWM, Sigma-Aldrich, UK) or the same volume of PBS into 15ml sterile tissue culture tubes (Fisher). Following incubation at 37°C for 48 hours, samples were centrifuged at 300 x *g* for 5min. Supernatants were collected and stored at −20°C until assayed for cytokine production.

Capture ELISAs were performed to quantify the secretion of selected cytokines representing different Th subsets: interferon (IFN)-γ (Th1), interleukin (IL)-4 (Th2), IL-17A (Th17) and IL-10 (Treg), following stimulation with PWM. All incubations were carried out at room temperature unless stated otherwise. IL-4 and IFN-γ were quantified using commercial ELISA kits according to the manufacturer’s instructions (MABTECH AB, Augustendalsvägen, SE, Sweden). For the quantification of IL-17A, polyclonal rabbit anti-bovine IL-17A antibodies were used alongside bovine recombinant protein (Kingfisher Biotech, Inc., St. Paul, MN). Mouse monoclonal anti-bovine IL-10 capture and detection antibodies (clones CC318 and CC320b respectively, BioRad) and standard curves produced using supernatants from COS-7 cells transfected with bovine IL-10 (Kwong *et al.* 2002; Corripio-Miyar *et al.* 2015) were used to quantify IL-10 secretion. Washing steps for all ELISAs were performed 6 times with 350μl washing buffer (Phosphate Buffered Saline (PBS) + 0.05% Tween 20) using a Thermo Scientific Wellwash™ Versa (ThermoFisher). High-binding capacity ELISA plates (Immunolon™ 2HB 96-well microtiter plates, ThermoFisher) were incubated with coating antibodies overnight at 4°C. Plates were then washed and blocked for 1h with PBS containing 0.05% Tween 20 (Sigma, UK) and 0.1% BSA Bovine Serum Albumin (BSA, Sigma, UK) for IL-4, IFN-γ and IL-17A or PBS containing 3% of BSA for IL-10. Following a further washing step, 50μl of supernatants or standards were added in duplicate for 1h. Subsequently, plates were washed and detection antibodies added for 1 h. This was followed by washing and addition of Streptavidin-HRP (Dako, Agilent, Santa Clara, US) for 45 min. After the final washing step, SureBlue TMB substrate (Insight Biotechnology, London, UK) was added and the reaction was stopped by the addition of TMB stop solution (Insight Biotechnology, London, UK). Absorbance values were read at O.D. 450nm. In order to quantify the cytokines of interest, samples were analysed 1:20, 1:4, neat or 1:4 for IFN-γ, IL-4, IL-17A and IL-10 respectively. Standard curves were included in all plates and were constructed using 7 serial dilutions of recombinant cytokines ranging from 6.25 to 400 pg/ml for IFN-γ (MABTECH AB); 31.25 to 2,000 pg/ml for IL-4 (MABTECH AB); 23.43 to 1,500 pg/ml for IL-17A (Kingfisher) and 0.206 to 13.2 BU/ml for IL-10 (Kwong *et al.* 2002). Finally, and in order to remove for the non-specific, natural cytokine release in all samples, results from the PWM stimulated samples were expressed as the corrected value of the cytokine release by subtracting the value obtained from PBS samples (background control) and multiplying by their corresponding dilution factor.

### Flow cytometry analysis

To quantify the leukocytes present in each blood sample, following gentle mixing of heparinised blood, leukocytes were enumerated using a Nucleocounter NC-200 (ChemoMetec, Denmark). Briefly, 20μl of heparinised blood were mixed with 180μl of Solution 17 (ChemoMetec) and heated at 37°C for 10min. The sample was then mixed to obtain a homogenous suspension and loaded into a Via1-Cassette™ (ChemoMetec) prior to counting on a Nucleocounter NC-200 cell counter. Total leukocyte counts per ml blood were recorded for each blood sample and used for the calculation of total cell counts expressing each of the transcription factors investigated.

Multiple colour flow cytometric analysis was carried out on blood samples from 188 individuals in order to identify and quantify the CD4 T helper cells expressing each the four master transcription factors (i.e., Tbet for Th1, Gata3 for Th2, RORγt for Th17, and FoxP3 for Treg).. Briefly, an aliquot of 2ml of blood was incubated with 10ml of warm red blood cell (RBC) lysis buffer (1.5M NH4Cl, 100mM NaHCO3, 10mM N2 EDTA in ddH_2_O) for 2min or until lysis was complete. Following two washes with PBS, Zombie Violet™ Fixable dead cell stain (Biolegend, US) was added to all samples and incubated for 15min at RT in the dark. Cells were then washed with PBS at 300 x *g* for 5min and stained with the cell surface antibody CD4 labelled to Alexa Fluor® 647 at pre-optimised concentrations for 20min at RT in the dark (see Table S1 for antibody details). Cells were then washed twice with FACS buffer (PBS + 5%FBS + 0.05%NaN3) and fixed with Fix/Perm buffer for 30min at 4°C (FoxP3 Staining Buffer Set buffer, Miltenyi Biotec, Bergisch Gladbach, Germany) according to manufacturer’s protocol. Following two washes with FACS buffer, cells were re-suspended in 1ml of PBS and stored at 4°C. Samples were then transferred to Moredun Research Institute (MRI) and analysed for the expression of Th-specific transcription factors within a month of sample collection. Briefly, following permeabilisation, monoclonal antibodies specific for the following Th-associated transcription factors Tbet (Th1), Gata3 (Th2), RORγt (Th17), and FoxP3 (Treg) alongside Isotype control antibody, all conjugated to phycoerythrin (PE), were added to samples and incubated for 30min in the dark at 4°C (see Table S1 for antibody details). Following staining, cells were washed twice with Permeabilization buffer and analysed immediately. A minimum of 100,000 events were acquired using a Sony SA3800 Spectral Analyzer (Sony Biotechnology, Ltd) and analysed using FlowJo vX for Windows 7.

In order to calculate the percentage of CD4^+^ and T cells expressing each of the hallmark Th-associated transcription factors (Figure 1), the following gating strategy was carried out. Following dead cell and doublet discrimination (Figure 2A&B), a leukocyte gate was created which included all white blood cells present (peripheral blood mononuclear cells (PBMC), granulocytes and neutrophils, Figure 2C). PBMC were then gated based on the forward scatter (FSC-A) and side scatter (SSC-A) (Figure 2D) and a CD4 gate created based on the fluorescence minus one (FMO) controls (Figure 2E). Threshold levels which determined positivity of each of the transcription factors in CD4^+^ T cells were also set using FMO controls. Consequently, the data obtained represented the percentage of CD4^+^ T cells in PBMC (Figure 2F) and the percentage of CD4^+^ T cells expressing Tbet, Gata3, RORγt or FoxP3 (Figure 2G-J). These were then expressed as a percentage of the total leukocyte population, and total blood leukocyte count data collected in the field was used to calculate the total numbers of CD4^+^ cells expressing each transcription factor per ml of blood.

**Figure 2.**
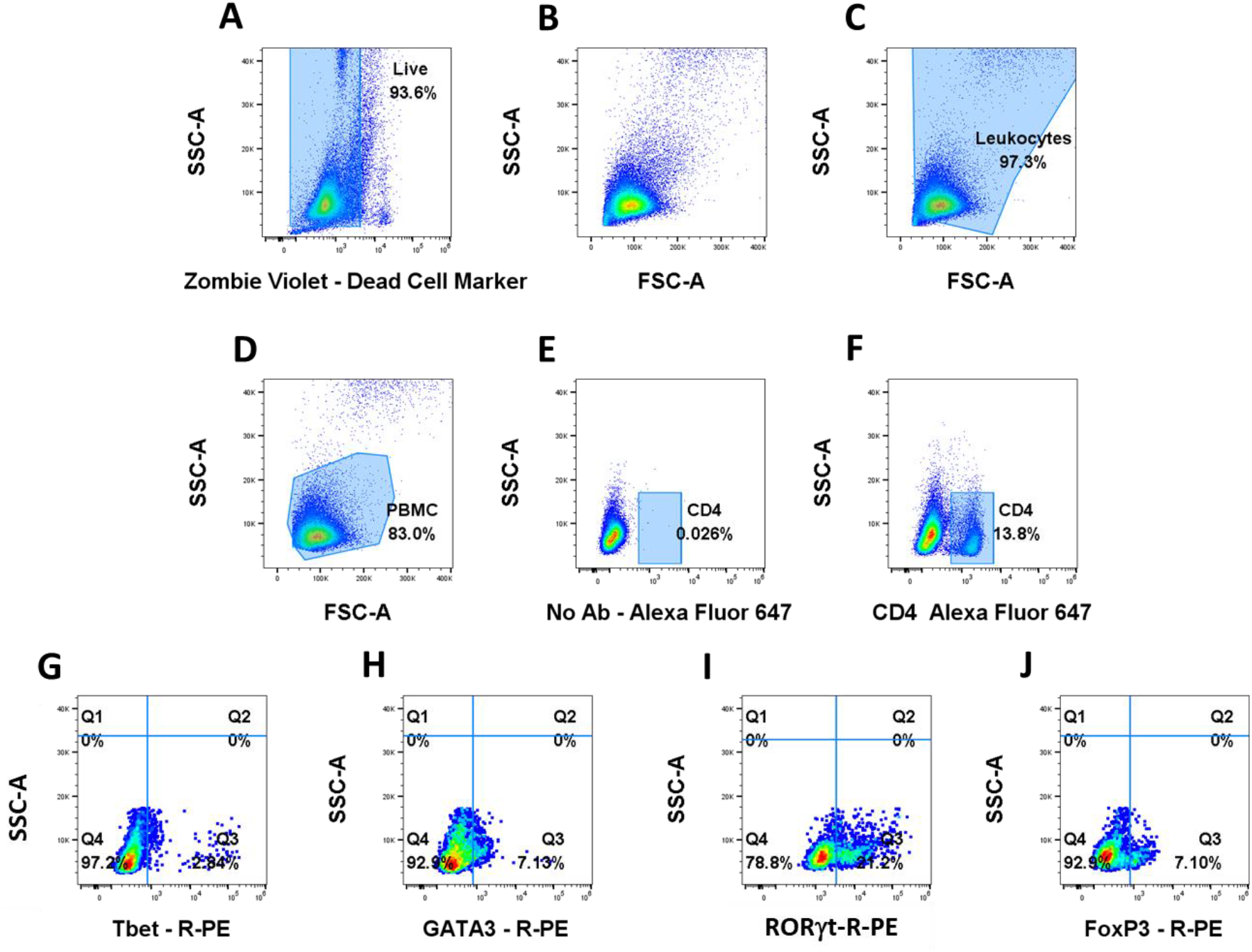
Gating strategy in flow cytometry analysis. The expression of CD4 and the transcription factors expressed by CD4^+^ T cells was studied by multi-colour flow cytometry. Cells were gated to eliminate dead cells (A) and doublets (B). Leukocytes were gated (C) followed by PBMC (D). Expression of CD4 conjugated to Alexa Fluor® 647 (F) in PBMC was determined by the gates set on no antibody controls (E). Gates for expression of transcription factors Tbet (G), Gata3 (H), RORγt (I) and FoxP3 (J) all conjugated to R-PE by CD4 were set using FMO controls. Data shown is from one representative individual with plots G-J corresponding to the expression of transcription factors by CD4^+^ T cells. A minimum of 100,000 events were acquired.

### Faecal egg/oocyst counts

Faecal samples were collected rectally and faecal egg counts (FEC) for strongyle nematodes and faecal oocyst counts (FOC) for coccidia were conducted on 2g samples using a modified salt-flotation technique (Jackson 1974) which is able to estimate FEC/FOC down to a resolution of 1 egg/oocyst per gram of faeces. These counts represent the most common parasites found in the Soay sheep in St Kilda, with strongyles comprising gastrointestinal nematode species (*Teladorsagia circumcincta, Trichostrongylus axei, Trichostrongylus vitrinus, Chabertia ovina, Bunostomum trigonocephalum* and *Strongyloides papillosus*) and coccidia comprising 11 *Eimeria* species (Wilson *et al.* 2004; Craig *et al.* 2006). Samples were stored anaerobically at 4°C until processed in our laboratory at MRI, around 2 weeks post collection. Briefly, 2g of faecal material was homogenised with 20ml of tap water. A subsample of 10ml was filtered through a sieve into a beaker and washed with 5ml of fresh water. Following transferring of the filtrate to a 15ml tube, samples were centrifuged at 200 x *g* for 2min. Supernatant was discarded, pellets resuspended into 10ml of saturated NaCl solution and centrifuged at 200 x *g* for 2min. Subsequently, tubes were clamped below the meniscus using forceps and parasite eggs/oocysts present in the surface of the saturated NaCl solution were transferred into a cuvette, filled with NaCl solution and parasite eggs/oocysts counted under a microscope.

### Statistical analysis

Although we collected blood samples from 238 sheep in total, due to time constraints in the field we were only able to prepare samples for 211 cytokine assays (IFN-γ, IL-4, IL-17A and IL-10) and 188 flow cytometry assays (total PBMCs, CD4^+^, CD4^+^Tbet^+^, CD4^+^Gata3^+^, CD4^+^RORγt^+^ and CD4^+^FoxP3^+^ cells). Of the 188 flow cytometry-prepared samples, 2 were not prepared for cytokine assays, leaving a total of 186 samples with all ten immune measures available. Faecal samples for parasite counts (FEC and FOC) were obtained for 229 of the 238 sampled sheep. Of the 186 fully immunologically sampled sheep, five were not faecal sampled, leaving 181 sheep with full immunological and parasitological data available (Table S2).

All analyses were conducted in R ver 3.6.3 (R Development Core Team 2019). First, we assessed the correlations among each of the 10 immune parameters that we measured. To ensure that any observed correlations were not due to common age- and sex-related variation among variables, we corrected for age and sex by fitting (Generalised) Linear Models (GLMs) for each variable with sex and age as an 11-level factor (ages 0-10) and their interaction. We assessed the best error distribution for each of the 10 parameters: we fitted linear models to IFN-γ, IL-10 and IL-17A; we fitted linear models to log-transformed IL-4; and fitted negative binomial generalised linear models (GLM) using the “MASS” package (Venables & Ripley 2002) for the other variables. We used the residuals from these models in our correlation analysis to present age- and sex-corrected results. We estimated Spearman’s rank correlations among all immune measures, assessing statistical significance using the “cor.mtest” function in the R package “corrplot” (Wei & Simko 2017).

We also explored the dimensionality of the raw (uncorrected) data using principal components analysis (PCA) with the function “prcomp”. Finally, we used the “adonis” function in the package “vegan” (Oksanen et al 2019) to run a PERMANOVA analysis in order to test for variation between sexes and age categories in distance matrices of the 10 uncorrected immune variables. Age was fitted as a four-level categorical variable, with lambs (aged ~4 months), yearlings (aged ~16 months), adults (aged 2-6 years) and geriatrics (aged 7+).

We next explored variation in each of the 10 immunological variables in related to sex and age (see Table S2 for sample sizes per immune measure). Since there were very few animals of very old age, females aged 10 and over were assigned age 10 and males aged 7 and over were assigned age 7. For cell phenotype traits (PBMC, CD4^+^, Tbet^+^, Gata3^+^, RORγt^+^ and FoxP3^+^ cells) we used a data set of 188 animals where information on all these traits, plus age and sex were available (Table S2); for the cytokines, we used a data set of 208 animals (of the 211 animals with cytokine data, 3 were of unknown age; Table S2). For each trait, we ran 14 different models with age categorised in different ways in order to best capture age- and sex-specific variation. Error structures were as given above, and as shown in Table S4. As well as a null model (0) and a model with sex only (1), we ran models (2-5) with linear age, quadratic age, age as a two-level factor (lambs versus others) and age as a four level factor (as described above), respectively. We also ran models 2-5 with the added effect of sex (models 6-9) and models 6-9 with the interaction between age and sex (models 10-13). Models for each trait were compared with Akaike Information Criterion (AIC) values, where the lowest value was considered to best fit the data, unless a simpler model had ΔAIC ≤ 2, in which case we selected the simpler model.

Finally, we tested for associations of each of the 10 immune parameters with strongyle FEC and coccidian FOC using negative binomial GLMs. For cell phenotype traits we used a data set of 183 animals and for the cytokines, we used the data set of 203 animals – these are the same as the data sets used for the analysis with regard to age and sex, minus five animals with missing parasitology data (Table S2). We included sex, age as a two-level factor (lambs v others), and their interaction in all models. We first fitted each of the 10 immune parameters in turn to models of FEC and FOC, and assessed their statistical significance against a model omitting the immune variables using likelihood ratio tests (LRTs). Next, we fitted interactions between the immune parameters and our two-level age category and assessed significance with LRTs, in order to test for differences between lambs and adults in how immune responses were associated with FEC. Finally, where more than one term was shown to be supported, we fitted all supported immune variables and their interactions into the same GLM in order to determine their independent associations with FEC or FOC.

## RESULTS

The correlations between the ten immune variables, after accounting for age and sex differences, are summarised in Figure 3. The correlations were mostly positive, with the strongest associations observed between the different cell counts and between IFN-γ and IL-4. In general, cytokines were only weakly associated with cell counts. To explore the possibility that the strong positive associations between cell phenotypes was driven by variation in total PBMC numbers, we re-ran the correlation analysis with the proportion of cells of each type within the total leukocyte population rather than the absolute numbers of cells. Since some cells don’t express any of the measured markers, these proportions would not necessarily be negatively associated. We found that, in general, associations between cell phenotypes expressed as proportions were relatively weak, suggesting that indeed the strong positive associations were driven by variation in cell numbers (Figure S1).

**Figure 3.**
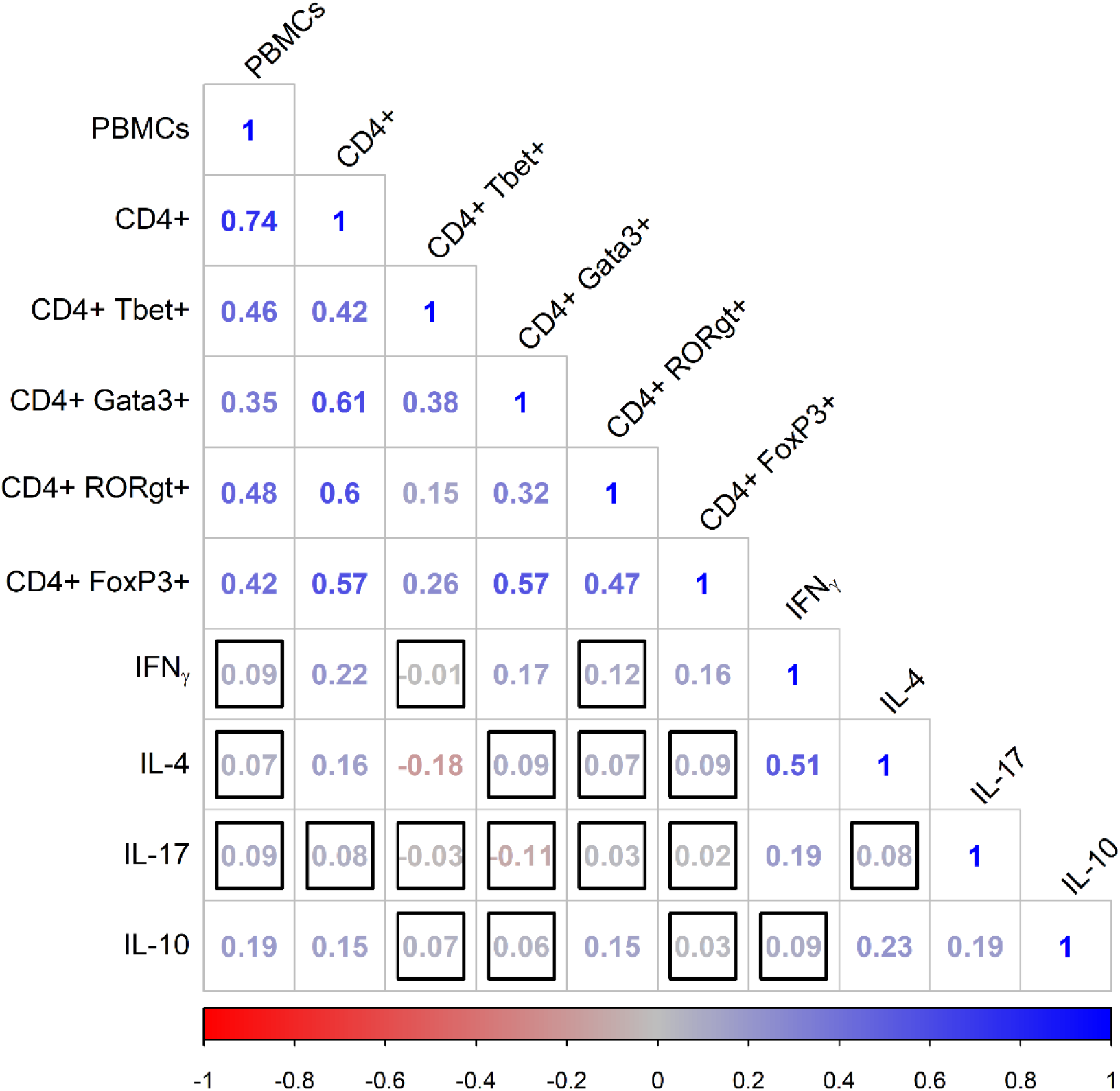
Correlation matrix showing Spearman’s rank correlations between pairs of immunological variables corrected for age and sex, where redder values indicate increasingly negative associations and bluer values indicate increasingly positive associations. Cell phenotypes represent cell counts per ml blood. Correlations outlined with a black square were not statistically significant at α=0.05.

The first principal component of our immune variables explained 32% of the variation and the second explained 25%; the subsequent axes explained 11% or less (Table S3). The first axis had strong negative loadings with all of the cell count variables, while the second axis had strong negative loadings with all four cytokines (Figure S2). Plotting our data on the first two principal components (PCs) revealed little evidence of differentiation between the sexes based on the first two PCs, but small differences with age were apparent (Figure S2). Specifically, older animals had lower values of PC2, suggesting stronger cytokine responses with age. This multivariate variation with age but not sex was supported by PERMANOVA analysis: while there was no evidence for an interaction between age category and sex (*F_DF=3_* = 0.34, P = 0.901) or the main effect of sex (*F_DF=1_* = 0.47, P = 0.592), there was some evidence for variation with age category (*F_DF=3_* = 3.54, P = 0.006). Age category, however, only explained a small proportion of the overall variation (R^2^ = 0.06).

This pattern of sex- and age-specific variation was also apparent in our individual analyses of each immunological variable (Tables S4 & S5). Variation in the number of PBMC and CD4^+^ cells was best explained by age as a four-level factor and both appeared to be at their lowest levels in adults compared to the other age groups (Figure 4A&B). The best-supported model for CD4^+^Tbet^+^ cells suggested higher cell numbers in lambs than in other age classes (Figure 4C), but there was no evidence for variation with either age or sex in CD4^+^ cells expressing the other three transcription factors (Figure 4D-F). Finally, all of the cytokines varied with age and/or sex, with a broad trend for increases with age and, where there were sex differences, greater increases in males than females. IFN-γ followed a quadratic trajectory with age, with a steep increase from younger ages to around age four, followed by a shallower increase thereafter (Figure 4G). IL-4 followed a similar pattern, although the best-supported model had age as a factor with four levels: while IL-4 was low in lambs, it increased dramatically in yearlings, with subsequently smaller increases in adults and then geriatrics (Figure 4H). The best-supported models for both IL-17A and IL-10 suggested an interaction between age and sex, with linear increases with age in both sexes, but a particularly pronounced increase in males compared to females (Figure 4I&J).

**Figure 4.**
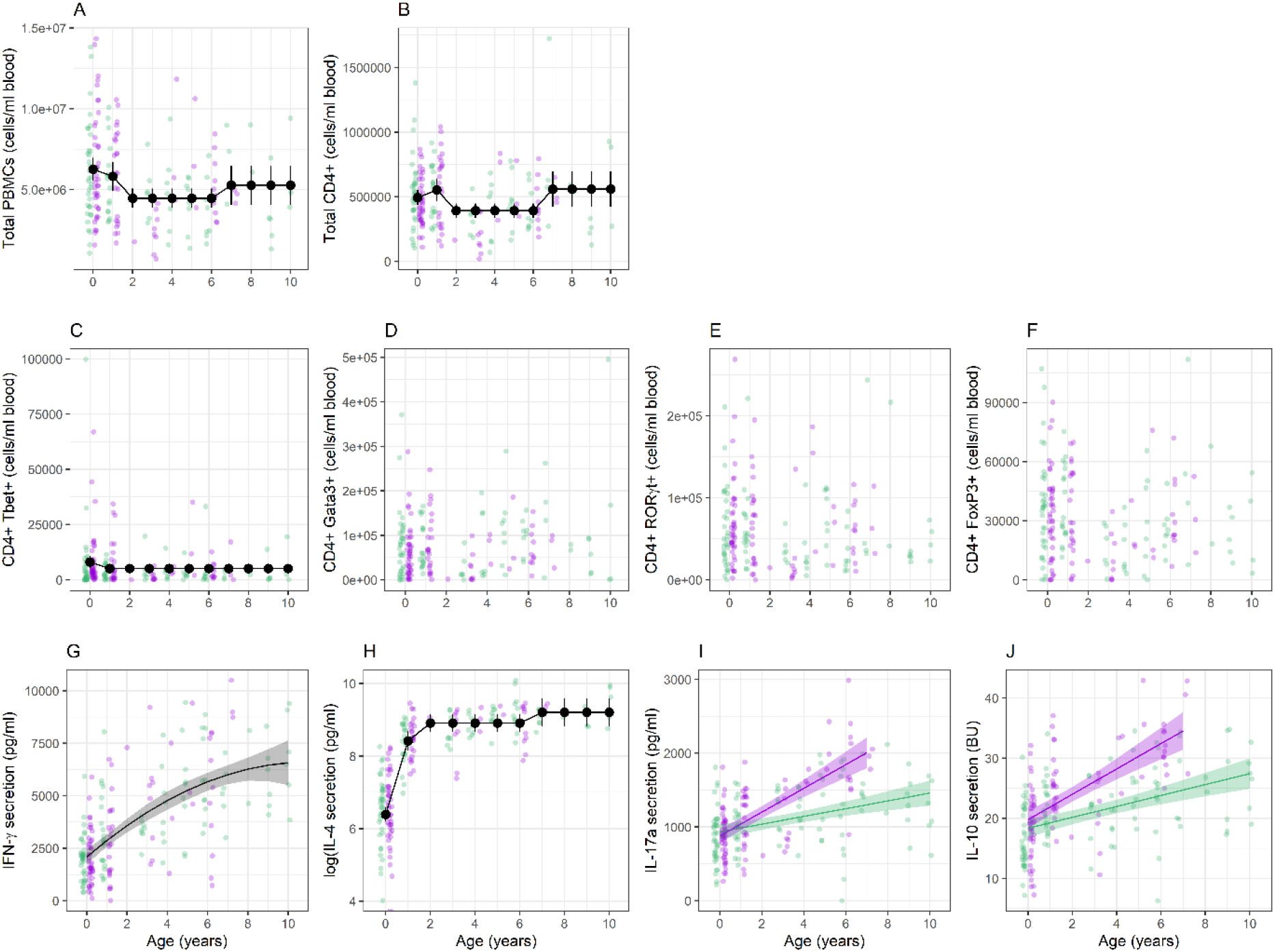
Age- and/or sex-specific variation in (A) PMBC; (B) CD4^+^ cells; (C) CD4^+^Tbet^+^ cells; (D) CD4^+^Gata3^+^ cells; (E) CD4^+^RORγt^+^ cells; (F) CD4^+^FoxP3^+^ cells; (G) IFN-γ; (H) IL-4; (I) IL-17A; and (J) IL-10. Points show raw data, with green representing females and purple showing males. Large points connected by lines show estimates ±95%CI from a model where age was fitted as a factor (with 2 or 4 levels) and lines with shaded areas show estimates ±95%CI from models where age was fitted as a continuous variable; green lines represent females and purple lines males. For model details, see Tables S4&S5.

Two of our immunological variables were associated with stongyle FEC as main effects: CD4^+^Gata3^+^ cells and IL-4 (Table 1; Figure 5B&E). Meanwhile, total PBMC, CD4^+^RORγt^+^ and CD4^+^FoxP3^+^ cells were all associated with FEC in interaction with age class as a two-level factor (Table 1). In addition, the interaction between age and CD4^+^Tbet^+^ cells was marginally non-supported (Table 1). In all cases, higher values were associated with lower FEC in lambs, but not in adults (Figure 5A; C&D). When we fitted all five of these supported terms (CD4^+^Gata3^+^ + IL-4 + Age*PBMC + Age*CD4^+^RORγt^+^ + Age*CD4^+^FoxP3^+^) into the same model, the main effects of CD4^+^Gata3^+^ and IL-4, and the interaction between age and CD4^+^FoxP3^+^ were still supported, while interactions between age and PBMCs and CD4^+^RORγt^+^ was not (Table S6). Only two of our immunological variables were associated with coccidian FOC (Table 1): both IFN-γ and IL-4 were negatively associated with FEC (Figure 6). However, when we fitted both of these into the same model, IFN-γ was still supported (estimate = −1.26E-04±4.94E-05, χ^2^ = 6.03, P = 0.014) but IL-4 was not (estimate = 1.03E-05±3.16E-05, χ2 = 0.12, P = 0.727).

**Figure 5.**
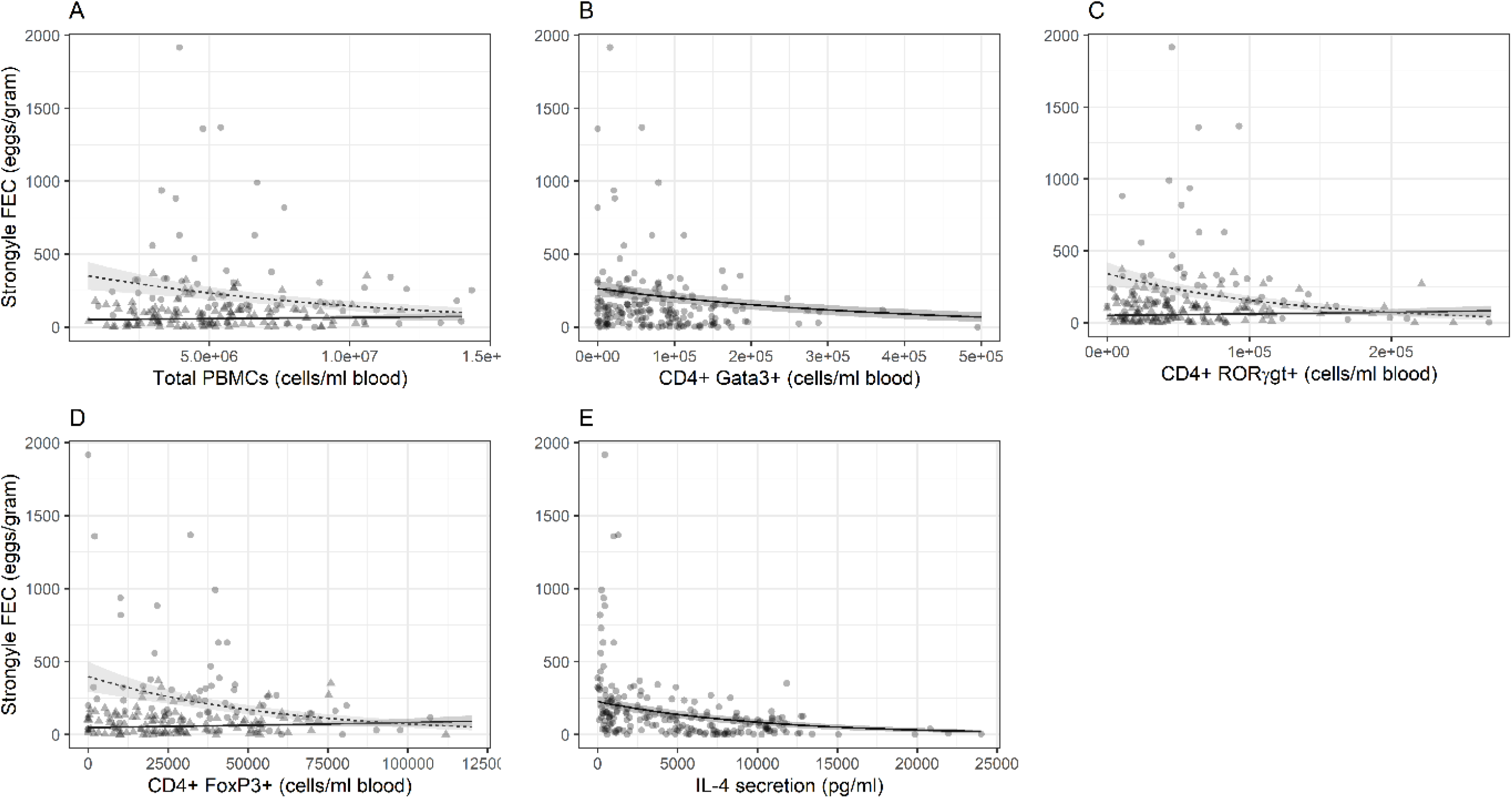
Associations between strongyle faecal egg count (FEC) and (A) total PBMCs; (B) total CD4^+^Gata3^+^ cells; (C) total CD4^+^RORyt^+^ cells; (D) total CD4^+^Foxp3^+^ cells; (E) IL-4 secretion. Points show raw data and lines show predictions from models in Table 1. Lambs are represented by circles and broken lines, adults by triangles and broken lines.

**Figure 6.**
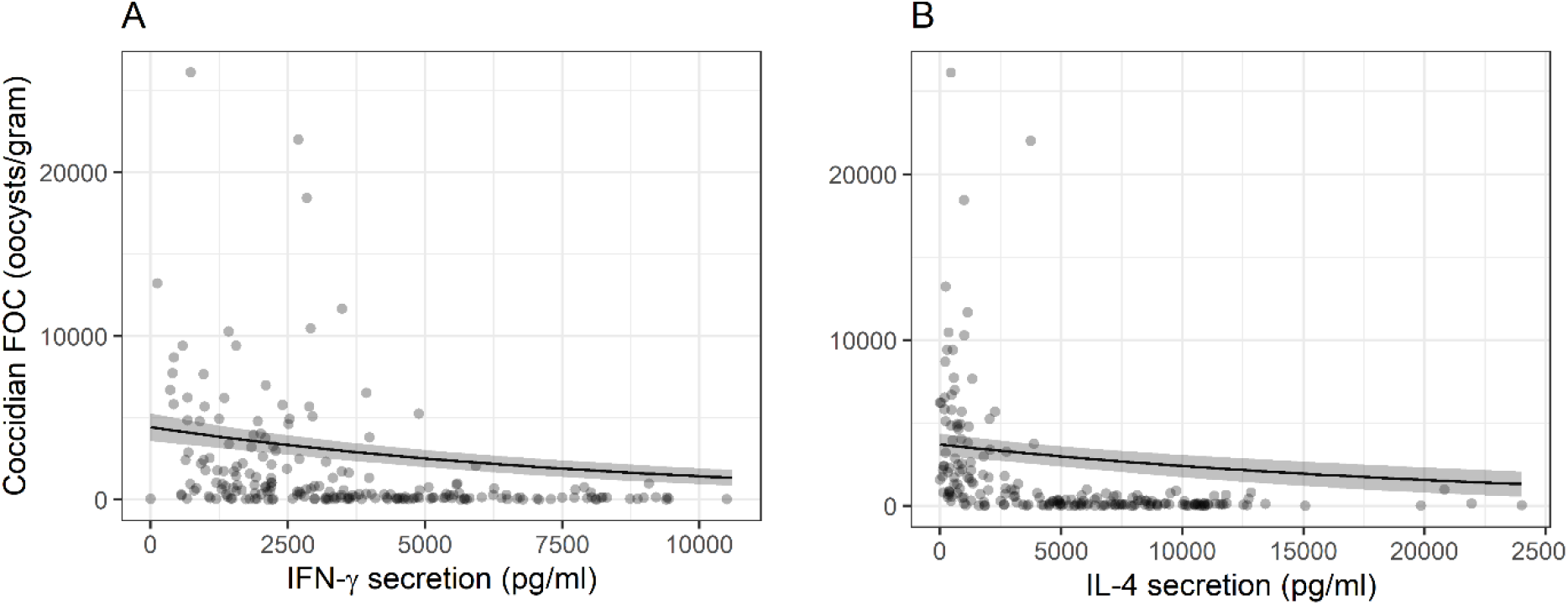
Associations between coccidian faecal oocyst count (FOC) and (A) IFN-γ secretion and (B) IL-4 secretion. Points show raw data and lines show predictions from models in Table 1.

**Table 1.**
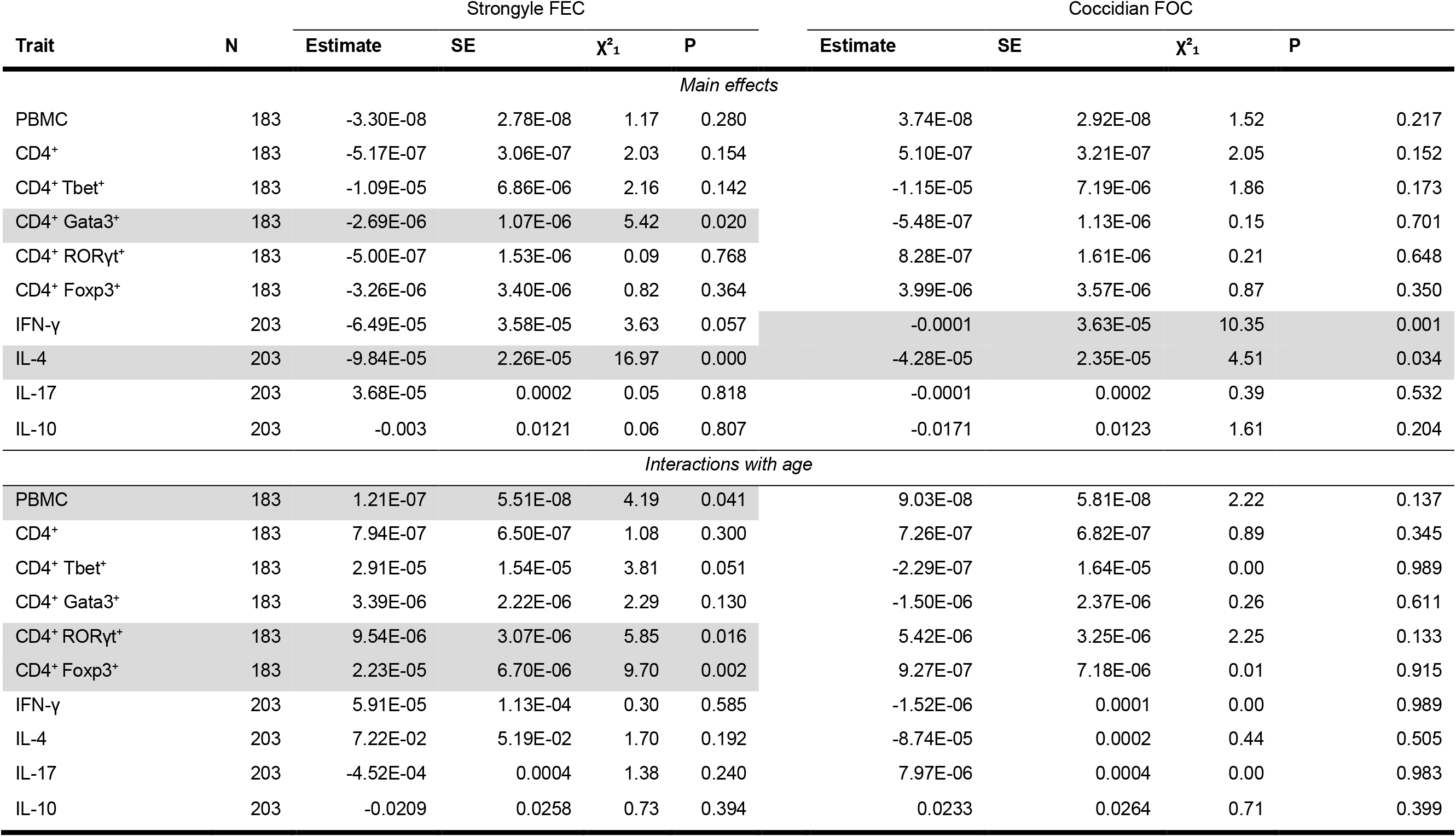
Results of generalised linear model analysis of associations between either strongyle faecal egg count (FEC) or coccidian faecal oocyst count (FOC) and each of our immunological parameters. Estimates and test statistics are shown for models where only the variable indicted with included in the model; interaction estimates show difference in slope between lambs and adults. Associations significant at α=0.05 are highlighted in grey.

## DISCUSSION

The adaptive immune system is critical to effective, long-lasting ability to respond to infection in vertebrates and research in medical, veterinary and laboratory animal settings illustrate the key role that T-helper (Th) cells play in orchestrating the adaptive response (Ovington, Alleva & Kerr 1995; Gill *et al.* 2000; Abolins *et al.* 2017). Nevertheless, measuring the Th response in naturally-infected populations will facilitate a more detailed understanding of the mechanisms underpinning variation in host responses to infection in food-limited, outbred populations, and enable analysis of how natural selection has shaped variation in immune responses (Pedersen & Babayan 2011; Maizels & Nussey 2013). In this study, we characterised variation in Th responses in a wild population of Soay sheep by enumerating cells expressing Th-specific transcription factors and measuring levels of canonical Th cytokines following *ex vivo* T cell stimulation. Our results highlight the importance of *how* we measure T helper responses, suggesting *ex vivo* stimulation assays may provide more ecologically-relevant assays of functional immune responses. Th cell counts and *ex vivo* cytokine production were weakly correlated, with only the latter showing the expected patterns of variation with age and independently predicting both strongyle and coccidian parasite burdens. We found support for the predictions that raised Th1 responsiveness (IFN-γ) would negatively associate with coccidian oocyst counts, while Th2 responsiveness (IL-4) would negatively associate with strongyle egg counts. However, contrary to our predictions, we found positive rather than negative associations between Th1 and Th2-associated measures. This provides rare support from outside the laboratory for the importance of different Th subsets for resistance to different kinds of parasite, but adds to mounting evidence from natural systems that Th1/Th2 trade-offs observed in reductionist laboratory experiments may not readily translate to complex natural systems (Arriero *et al.* 2017; Young *et al.* 2020).

### Variation in T helper phenotype with age and sex

Age-specific variation in immunity is widely observed and expected: the immaturity of the immune system in juveniles is often associated with higher parasite burden and less effective responses (Woolhouse 1998) and this is certainly also true of domestic sheep infected with strongyle nematodes (Gibson & Parfitt 1972; Smith *et al.* 1985). In later adulthood, immunosenescence is often detected in wild vertebrates, particularly in adaptive responses (Peters *et al.* 2019). We have previously observed increases in circulating strongyle-specific antibody levels between lambs and adults in the Soay sheep (Sparks *et al.* 2018) and senescent declines in later life that are associated with increased risk of mortality (Froy *et al.* 2019). We have also observed pronounced declines with age in the proportions of certain T cell sub-types in separate cross-sectional studies of the Soay sheep, most notably naïve (CD45RA^+^) helper (CD4^+^) and cytotoxic (CD8^+^) T cells and γδ^+^ T cells (Nussey et al 2012, Watson et al 2016). However, these changes were very much expected based on fundamental processes in immune development, such as thymic involution, and our previous studies did not tease apart different functional T helper subsets (Nussey *et al.* 2012; Watson *et al.* 2016). Here, we found little evidence of age-dependence in the number of cells expressing transcription factors associated with different T helper subsets (Figure 4). However, we did find pronounced increases from lambs to adults in *ex vivo* T cell cytokine responses associated with all Th subsets. One possible explanation for this pattern is that, as animals age and are exposed to parasite antigens, there is an expansion of Th memory pools which have a faster and more rapid cytokine response following activation (Pennock *et al.* 2013), potentially explaining why Th cell counts are static but cytokine responses increase with age. However, regardless of the underlying mechanism, our results suggest that variation in the number of functionally distinct Th cells, within the circulatory pool at least, does not play a significant role in the well-documented age-related variation in immunity in our study system.

Increases in cytokine responses with age have been previously described in healthy human populations (Chipeta *et al.* 1998), where a Th1 polarisation of cytokine responses can be observed with age (Krampera *et al.* 1999; Sakata-Kaneko *et al.* 2000). We observed no signs of such a polarisation: production of cytokines associated with Th1, Th2, Th17 and Treg functional responses all increased with age. However, most human studies focus on cytokine production specifically by CD4^+^ and CD8^+^ cells, whereas our assays include cytokine production by all lymphocytes. It is known that activated B cells and natural killer cells can produce IL-10 and IFN-γ, respectively, and may be contributing to the cytokine responses we measured (Gray & Horwitz 1995; Varma *et al.* 2002; Duddy, Alter & Bar-Or 2004). Although a more focused study of the cytokines produced by CD4^+^ T cells might have shown different results, this lack of a shift towards Th1 responses with age could also reflect the consistent life-long exposure to gastrointestinal parasites experienced by wild Soay sheep. This could produce a more pronounced increase in other functional subsets associated with mucosal immune responses with age (e.g. Th2 and Th17) than observed in lab rodents and Western humans. Furthermore, gastrointestinal nematodes are known to produce and induce regulatory immune responses in sheep (McNeilly *et al.* 2013), and this could contribute to increases in IL-10 responsiveness with age.

We found little evidence for sex differences in Th phenotypes in wild Soay sheep. Sex-specific variation in defence against infection is predicted and has been routinely observed in wild populations (Restif & Amos 2010). Males often exhibit increased parasite burden and less effective immune responses than females, and this has been attributed to various causes, including behaviour, resource allocation, and the putative immunosuppressive effects of testosterone (Foo *et al.* 2017). Previous work on the Soay sheep has shown sex differences in anti-strongyle antibody levels, with males showing weaker responses (Hayward et al 2014, Sparks et al 2019) and higher burdens of nematode parasites across age groups (Craig *et al.* 2006; Hayward *et al.* 2009). In other wild mammals, male voles exhibited lower levels of expression of the Th2-associated transcription factor Gata3 in peripheral blood and wild male badgers showed lower IFN-γ responses than females following stimulation with PWM, although no sex differences were found in wild buffalo (Ezenwa *et al.* 2010; Beirne *et al.* 2016; Arriero *et al.* 2017). Generally, our data provide limited evidence for an important role of T helper cell responsiveness underpinning the widely observed sex differences in parasitism and immune responsiveness in wild vertebrates. However, we note that *ex vivo* production of Th17 and Treg cytokines did increase more rapidly with age in males than females, and thus may warrant further investigation, and that our relatively small, cross-sectional data set may have prevented us from detecting more subtle longitudinal changes with age and differences between the sexes.

### Correlations among immune measures

Evidence from laboratory immunology has supported antagonistic interactions between Th1 and Th2 responses, such that expression of one is associated with down-regulation of another (Mosmann & Coffman 1989; Kaiko *et al.* 2008); in hosts infected with a range of parasites, we may therefore expect to see hosts trading-off Th1 responses against microparasites with Th2 responses against macro parasites (Cox 2001). Further, we would expect that hosts exhibiting high counts of one type of Th cell would also show high secretion of the corresponding canonical Th cytokine. Despite this expectation, we found limited evidence for associations between Th cell counts and the cytokines associated with their corresponding functional response; for example, high CD4^+^Tbet^+^ cell numbers were not associated with higher levels of IFN-γ production following stimulation (Figures 1 & 3). This could suggest that the steady-state (cell counts) and responsiveness (cytokine production) of Th measures provide different information about an individual’s immune responsiveness and state. Since our assays estimate cytokine production across all leukocytes, rather than just Th cells, this could also suggest that non-Th cells play an important role in the *ex vivo* cytokine responses we measured. Moreover, our specific prediction of a negative correlation between Th1 and Th2 phenotypes was not apparent in our data; indeed, the strongest positive association between cytokine responses was between the Th1 and Th2 cytokines, IL-4 and IFN-γ. Antagonism between Th1 and Th2 responses are well-established in laboratory immunology (Seder & Paul 1994; Mosmann & Sad 1996), and supported by experimental studies of wild buffalo in which Th2-inducing helminth parasites were removed and increased IFN-γ levels (Ezenwa & Jolles 2015). However, data from unmanipulated wild rodent populations provides evidence for synergistic, rather than antagonistic, associations between Th1 and Th2 phenotypes (Arriero *et al.* 2017; Young *et al.* 2020), and indeed work in domestic sheep indicates a complex and temporarily regulated interplay between Th1 and Th2 immune responses is associated with gastro-intestinal parasite resistance (Hassan *et al.* 2011). Variation in resource acquisition, which is common in natural systems, resulting in individual differences in the ability to invest resources in immunity, could readily drive positive associations between different arms of the immune response even if functional trade-offs are present (Noordwijk & Jong 1986; Arriero *et al.* 2017). Additionally, persistent challenge from a variety of parasites, which is a feature of natural populations, and the need for considerable plasticity in responses, could also lead to selection for the ability to amount effective immune responses to different parasites. As such, trade-offs between different arms of the Th response could be masked by variation in resource acquisition, coinfection, and the need to adjust responses in responses to ecological and epidemiological conditions.

### T helper phenotypes and parasite burdens

In line with our predictions, levels of the Th2-associated cytokine IL-4 were negatively associated with strongyle faecal egg count, while the Th1-associated cytokine IFN-γ was negatively associated with coccidian oocyst count. This represents the clearest evidence to date supporting the paradigm that Th1 and Th2 responses have distinct roles in tackling microparasite and macroparasite infections, respectively, from a wild population. In addition, for strongyle parasites, both numbers of CD4 T cells expressing Gata3 or Foxp3 (i.e. Th-2 and Treg polarised cells) were also negatively associated with faecal egg count. While the association with Gata3 provides further evidence that Th2 immunity is important for control of strongyle parasites, the association between Treg numbers and strongyle faecal egg count, which was only evident in lambs, is more counterintuitive, given that Treg induction has been suggested as an immune evasion strategy for these parasites (Grainger *et al.* 2010). However, the relationship between Treg and helminth immunity is complex – in mice it has been shown that while too many Treg impair Th-2 immunity and lead to chronic infections, too few lead to dysregulated immune responses and increased worm burdens (Smith *et al.* 2016). Similar findings have also been reported in sheep, with resistance to strongyle parasites being associated with an early Treg-Th2 immune response during primary infection (Hassan *et al.* 2011). Together these suggest that an optimal balance between Th-2 and Treg responses is critical to induce effective anti-strongyle immunity, at least during initial exposure to the parasite.

While both cytokine production and cell numbers associated with parasite burdens, *ex vivo* cytokine responses of lymphocytes were generally better at predicting variation in measures of parasite burden than cell counts of Th subtypes. This might be expected given the key role cytokines play in orchestrating effectors of the response, such as antibodies and immune effector cells (Abbas, Murphy & Sher 1996). Furthermore, the fact that cytokine but not cell number measurements vary with age and sex suggests they may reflect more useful and informative markers of immune function that could be applied more widely in field studies of non-model systems. This is promising in terms of future studies of adaptive immunity in wild populations given that whole blood *ex vivo* cytokine release assays are generally more easily adapted to the field than flow cytometry-based assays that require immediate cell isolation and labelling. Consequently, our results highlight the urgent need for expanding the immunological toolbox for non-model systems, as antibodies for cytokines and other reagents are not always readily available for such species.

Previous studies of wild mammals have provided evidence in support of associations between Th phenotypes and measures of parasite burden. For example, in a wild badger population, “excretors” of bovine tuberculosis (bTB; *Mycobacterium bovis)* had lower mean IFN-γ responses than did bTB-negative individuals (Beirne *et al.* 2016). In wild buffalo, anthelminthic-treated individuals had stronger IFN-γ responses and while treatment was not associated with bTB status, treated animals were less likely to die from bTB, suggesting a protective role for IFN-γ in bTB infection (Ezenwa & Jolles 2015). Finally, expression of Gata3 was positively associated with an index of macroparasite burden in wild adult voles, but negatively correlated in juvenile voles; this was interpreted as a Th2-mediated switch from resistance to tolerance with age (Jackson *et al.* 2014). These results echo findings in domestic sheep where lines artificially selected to be more resistant to helminth infection show enhanced IL-4 (Th2) responses (Terefe *et al.* 2007; Gossner *et al.* 2013; Zaros *et al.* 2014; Wilkie *et al.* 2016). Relatively little is known about coccidian immunity in domestic sheep, although there is convincing evidence that natural infection drives expression of IFN-γ and other Th-1 associated cytokines such as IL-2 and TNF-α within the intestinal mucosa (Ozmen, Adanir & Haligur 2012). In wild Soay sheep, strongyle faecal egg count is positively associated with coccidian oocyst count counts (Craig *et al.* 2008), and coupled with the positive association between IL-4 and IFN-γ, the picture appears to be one of variation in the ability to mount responses to diverse parasite taxa, with individuals able to respond effectively to strongyles also being able to respond effectively to coccidia. Our previous work on Soay sheep clearly shows that functionally distinct immune phenotypes (antibodies of different isotypes) can be positively correlated but show different patterns of association with fitness (Sparks *et al.* 2018). A crucial next step for our understanding of natural selection on T helper phenotype is to relate the variation in different functional measures to fitness in wild populations, and determine whether associations are mediated by effects on parasite burden or individual condition, e.g. (Sparks *et al.* 2020).

### Conclusions

Overall, our results provide important evidence that measuring immune phenotypes associated with different functional arms of the T helper response can help us to predict the parasite burden and infection status of wild mammals. Whilst ours is not the first study to provide evidence linking Th1- or Th2-associated cytokine production with infection status in wild mammals, we have assessed T helper phenotype in an unusually comprehensive fashion using reagents developed for veterinary immunology. Our results provide, to our knowledge, the first evidence simultaneously linking raised Th1-associated responses to reduced micro-parasite burdens and raised Th2-associated responses to reduced macro-parasite burdens in the wild. Whilst this supports the paradigm that different T helper arms have effector functions that protect from particular parasite groups, our data challenge the idea that investment in one Th arm constrains investment in another in the wild. One limitation of our study is its cross-sectional nature, with larger-scale longitudinal data required to determine whether age-related variation is driven by within-individual change or selective effects and how such variation impacts on demography (Nussey *et al.* 2008). The small number of longitudinal studies measuring Th-related phenotypes in the wild suggest they are weakly to moderately repeatable across repeated measurements (Beirne, Delahay & Young 2015; Arriero *et al.* 2017). Further studies examining longitudinal changes across functional Th subsets and relating these to parasite pressure and demographic rates are likely to illuminate the causes and consequences of variation in Th function for natural populations.

## Acknowledgements

This work was funded by a large NERC grant (NE/R016801/1), and the long-term study on St Kilda was funded principally by responsive mode grants from NERC. We thank the National Trust for Scotland for their ongoing support of our work on St Kilda, and QinetiQ and Kilda Cruises for logistical support. We are very grateful to the 2019 August St Kilda Soay sheep project catch team, without whom the samples we used in this study could not have been collected, and especially to Hannah Lemon, Amy Sweeny and Sanjana Ravindran for their help processing samples in the field.

