## Supplementary Files for "Functionally distinct T-helper cell phenotypes predict resistance to different types of parasites in a wild mammal"

**Table S1** Monoclonal antibodies used for flow cytometric analysis.

| **Antigen** | **Antibody clone** | **Fluorochrome** | **Isotype** | **Source** |
| --- | --- | --- | --- | --- |
| **CD4** | 44.38 | Alexa Fluor^®^ 647 | IgG2a | BioRad |
| **Isotype control** | REA293 | PE | Recombinant  human IgG1 | Miltenyi Biotec |
| **Tbet** | REA102/4B10 | PE | Recombinant  human IgG1 | Miltenyi Biotec |
| **Gata3** | REA174/TWAJ | PE | Recombinant  human IgG1 | Miltenyi Biotec |
| **RORγt** | REA278/AFKJS-9 | PE | Recombinant  Human IgG1 | Miltenyi Biotec |
| **FoxP3** | FJK-16s | PE | Rat IgG2a | eBioscience |

***Table S2.*** Sample sizes used in analyses, broken down by *age and sex. “Number sampled” gives the total number of animals that we had any data on; “FEC” gives the number of individuals with known parasite egg/oocyst counts; “Cells” gives the number of individuals with cell phenotype data (PBMC, CD4^+^, CD4^+^Tbet^+^, CD4^+^Gata3^+^, CD4^+^RORγt^+^ and CD4^+^FoxP3^+^ cells); “Cytokines” gives the number of individuals with known IFN-γ, IL-4, IL-17A and IL-10 data; “All” gives the number of animals where age, sex, parasite counts, cell phenotype and cytokines were measured.*

| **Sex** | **Age** | **Number sampled** | **FEC** | **Cells** | **Cytokines** | **All** |
| --- | --- | --- | --- | --- | --- | --- |
| Female | 0 | 39 | 37 | 34 | 37 | 32 |
| Female | 1 | 23 | 22 | 18 | 21 | 17 |
| Female | 3 | 10 | 10 | 6 | 8 | 6 |
| Female | 4 | 10 | 10 | 7 | 8 | 7 |
| Female | 5 | 9 | 9 | 8 | 9 | 8 |
| Female | 6 | 13 | 13 | 8 | 11 | 8 |
| Female | 7 | 9 | 9 | 4 | 6 | 4 |
| Female | 8 | 2 | 2 | 2 | 2 | 2 |
| Female | 9 | 7 | 7 | 5 | 6 | 5 |
| Female | 10 | 8 | 8 | 4 | 6 | 4 |
| Female | Unknown | 4 | 4 | 0 | 3 | 0 |
| Male | 0 | 49 | 47 | 41 | 43 | 38 |
| Male | 1 | 24 | 24 | 23 | 23 | 22 |
| Male | 2 | 1 | 1 | 1 | 1 | 1 |
| Male | 3 | 11 | 11 | 8 | 8 | 8 |
| Male | 4 | 3 | 3 | 3 | 3 | 3 |
| Male | 5 | 3 | 3 | 3 | 3 | 3 |
| Male | 6 | 10 | 10 | 10 | 10 | 10 |
| Male | 7 | 3 | 3 | 3 | 3 | 3 |
| Female | All | 134 | 131 | 96 | 117 | 93 |
| Male | All | 104 | 102 | 92 | 94 | 88 |
|  | Known | 234 | 229 | 188 | 208 | 181 |
|  | Total | 238 | 233 | 188 | 211 | 181 |

***Table S3.*** *Results of principal components analysis (PCA) of 10 immunological variables, showing the standard deviation and proportion of total variation explained by each of the principal components.*

|  | **PC1** | **PC2** | **PC3** | **PC4** | **PC5** | **PC6** | **PC7** | **PC8** | **PC9** | **PC10** |
| --- | --- | --- | --- | --- | --- | --- | --- | --- | --- | --- |
| Standard deviation | 1.80 | 1.57 | 1.06 | 0.98 | 0.78 | 0.75 | 0.65 | 0.53 | 0.43 | 0.41 |
| Proportion of variance | 0.32 | 0.25 | 0.11 | 0.10 | 0.06 | 0.06 | 0.04 | 0.03 | 0.02 | 0.02 |
| Cumulative proportion | 0.32 | 0.57 | 0.68 | 0.78 | 0.84 | 0.90 | 0.94 | 0.97 | 0.98 | 1.00 |

***Table S4.*** *AIC values from models testing for age- and sex-specific variation in each of 10 immunological variables. Age(2) and Age(4) refer to age modelled as a categorical variable with 2 or 4 levels respectively. For each trait, the model highlighted in* ***bold*** *is the model with the lowest AIC; the model highlighted in grey is the chosen model based on AIC. Where a model simpler than the model with lowest AIC had ΔAIC<2 relative to the model with the lowest AIC, the simpler model was chosen to represent the data. Errors structures are NB = negative binomial; LN = log-normal; G = Gaussian.*

| **Variables** | **PBMCs** | **CD4+** | **CD4+Tbet+** | **CD4+Gata3+** | **CD4+RORγt+** | **CD4+FoxP3+** | **IFN-γ** | **IL-4** | **IL-17** | **IL-10** |
| --- | --- | --- | --- | --- | --- | --- | --- | --- | --- | --- |
| *Error structure* | *NB* | *NB* | *NB* | *NB* | *NB* | *NB* | *G* | *LN* | *G* | *G* |
| Null | 6091.66 | 5188.76 | 3518.67 | **4574.31** | **4520.40** | **4262.99** | 3827.57 | 765.25 | 3163.19 | 1410.14 |
| Sex | 6092.66 | 5190.29 | 3518.65 | 4574.58 | 4521.53 | 4264.91 | 3826.16 | 761.66 | 3164.67 | 1407.61 |
| Age | 6087.14 | 5190.53 | 3517.70 | 4574.88 | 4521.59 | 4264.27 | 3713.62 | 659.73 | 3115.54 | 1366.61 |
| Age² | 6085.12 | 5189.49 | 3518.66 | 4576.26 | 4523.46 | 4265.05 | **3710.29** | 621.45 | 3104.14 | 1360.00 |
| Age(2) | 6086.14 | 5190.60 | **3517.16** | 4575.94 | 4521.72 | 4264.09 | 3766.43 | 577.14 | 3126.99 | 1357.28 |
| Age(4) | **6083.53** | **5182.91** | 3519.90 | 4578.68 | 4523.19 | 4264.29 | 3715.47 | **568.92** | 3114.44 | 1356.44 |
| Sex+Age | 6088.91 | 5191.88 | 3518.52 | 4575.80 | 4523.01 | 4266.07 | 3715.08 | 661.73 | 3109.63 | 1349.24 |
| Sex+Age² | 6086.62 | 5191.12 | 3519.15 | 4577.31 | 4524.80 | 4266.95 | 3711.98 | 623.09 | 3099.34 | 1343.88 |
| Sex+Age(2) | 6087.58 | 5192.09 | 3517.28 | 4576.35 | 4522.98 | 4265.96 | 3767.46 | 577.38 | 3126.14 | 1346.46 |
| Sex+Age(4) | 6085.07 | 5184.41 | 3520.64 | 4579.36 | 4524.37 | 4266.21 | 3717.25 | 570.43 | 3109.88 | 1340.56 |
| Sex*Age | 6090.71 | 5193.85 | 3520.44 | 4577.46 | 4525.01 | 4268.04 | 3713.39 | 659.33 | 3087.56 | **1337.68** |
| Sex*Age² | 6088.36 | 5192.81 | 3521.08 | 4579.19 | 4526.79 | 4268.48 | 3712.79 | 624.81 | **3086.60** | 1337.88 |
| Sex*Age(2) | 6089.57 | 5193.69 | 3519.00 | 4578.34 | 4524.98 | 4267.91 | 3768.65 | 579.33 | 3126.40 | 1347.13 |
| Sex*Age(4) | 6090.63 | 5189.81 | 3525.27 | 4584.56 | 4530.33 | 4271.87 | 3713.02 | 575.07 | 3107.73 | 1339.83 |

***Table S5.*** *Parameter estimates from the best-supported model of age and sex for each of the 10 immunological variables (model highlighted in grey in Table S4). Age(2) and Age(4) refer to age modelled as a categorical variable with 2 or 4 levels respectively. Further details about the selected models can be found in Table S4.*

| **Variable** | **PBMCs** | **CD4+** | **CD4+ Tbet+** | **CD4+ Gata3+** | **CD4+ RORγt+** | **CD4+ FoxP3+** |
| --- | --- | --- | --- | --- | --- | --- |
| Intercept | 15.6487 (0.0575) | 13.1089 (0.0607) | 8.9944 (0.1858) | 11.2429 (0.0932) | 11.0259 (0.0652) | 10.3556 (0.0845) |
| Age(2) - lamb | NA | NA | 0.0000 (0.0000) | NA | NA | NA |
| Age(2) - adult | NA | NA | -0.4454 (0.2397) | NA | NA | NA |
| Age(4) - lamb | 0.0000 (0.0000) | 0.0000 (0.0000) | NA | NA | NA | NA |
| Age(4) - yearling | -0.0729 (0.0967) | 0.1143 (0.1021) | NA | NA | NA | NA |
| Age(4) - adult | -0.3366 (0.0888) | -0.2269 (0.0938) | NA | NA | NA | NA |
| Age(4) - geriatric | -0.1712 (0.1306) | 0.1260 (0.1380) | NA | NA | NA | NA |
|  | **IFN-γ** | **IL-4** | **IL-17** | **IL-10** |  |  |
| Intercept | 2092.51 (178.83) | 6.3955 (0.1047) | 934.01 (52.00) | 18.3442 (0.7748) |  |  |
| Sex - female | NA | NA | 0.00 (0.00) | 0.00 (0.00) |  |  |
| Sex - male | NA | NA | -55.35 (73.13) | 1.4443 (1.0896) |  |  |
| Age | 819.12 (132.85) | NA | 52.54 (11.45) | 0.9083 (0.1706) |  |  |
| Age^2 | -37.10 (16.08) | NA | NA | NA |  |  |
| Age(2) - lamb | NA | NA | NA | NA |  |  |
| Age(2) - adult | NA | NA | NA | NA |  |  |
| Age(4) - lamb | NA | 0.0000 (0.0000) | NA | NA |  |  |
| Age(4) - yearling | NA | 2.0255 (0.1758) | NA | NA |  |  |
| Age(4) - adult | NA | 2.5151 (0.1592) | NA | NA |  |  |
| Age(4) - geriatric | NA | 2.8090 (0.2216) | NA | NA |  |  |
| Sex*Age | NA | NA | 108.69 (21.73) | 11999 (0.3237) |  |  |

***Table S6.*** *Parameter estimates for a model containing all terms that were statistically supported explanatory variables of FEC when fitted individually (Table 1). Estimates are from this full model containing all of the listed terms, while test statistics and p-values are from likelihood ratio tests comparing the full model with a model with the term of interest omitted. Terms in bold are statistically supported independently of the other variables in the model.*

| **Variable** | **Estimate** | **SE** | **χ²₁** | **P** |
| --- | --- | --- | --- | --- |
| Intercept | 6.3860 | 0.3000 |  |  |
| Age (lamb) | 0.0000 | 0.0000 |  |  |
| Age (adult) | -1.8860 | 0.4208 |  |  |
| Sex (female) | 0.0000 | 0.0000 |  |  |
| Sex (male) | 0.3439 | 0.2326 |  |  |
| Age (adult): Sex (male) | 0.4834 | 0.2978 | 2.57 | 0.109 |
| PBMCs | -4.43E-08 | 4.08E-08 |  |  |
| CD4^+^RORgt^+^ | -2.27E-06 | 2.69E-06 |  |  |
| CD4^+^FoxP3^+^ | -7.69E-06 | 5.54E-06 |  |  |
| **CD4^+^Gata3^+^** | **-3.24E-06** | **1.21E-06** | **4.77** | **0.029** |
| **IL-4** | **-8.97E-05** | **2.27E-05** | **13.81** | **<0.001** |
| Age (adult): PBMCs | 3.59E-08 | 6.29E-08 | 0.33 | 0.566 |
| Age (adult): CD4^+^RORgt^+^ | 1.99E-06 | 3.60E-06 | 0.21 | 0.647 |
| **Age (adult): CD4^+^FoxP3^+^** | **2.33E-05** | **7.79E-06** | **7.29** | **0.006** |

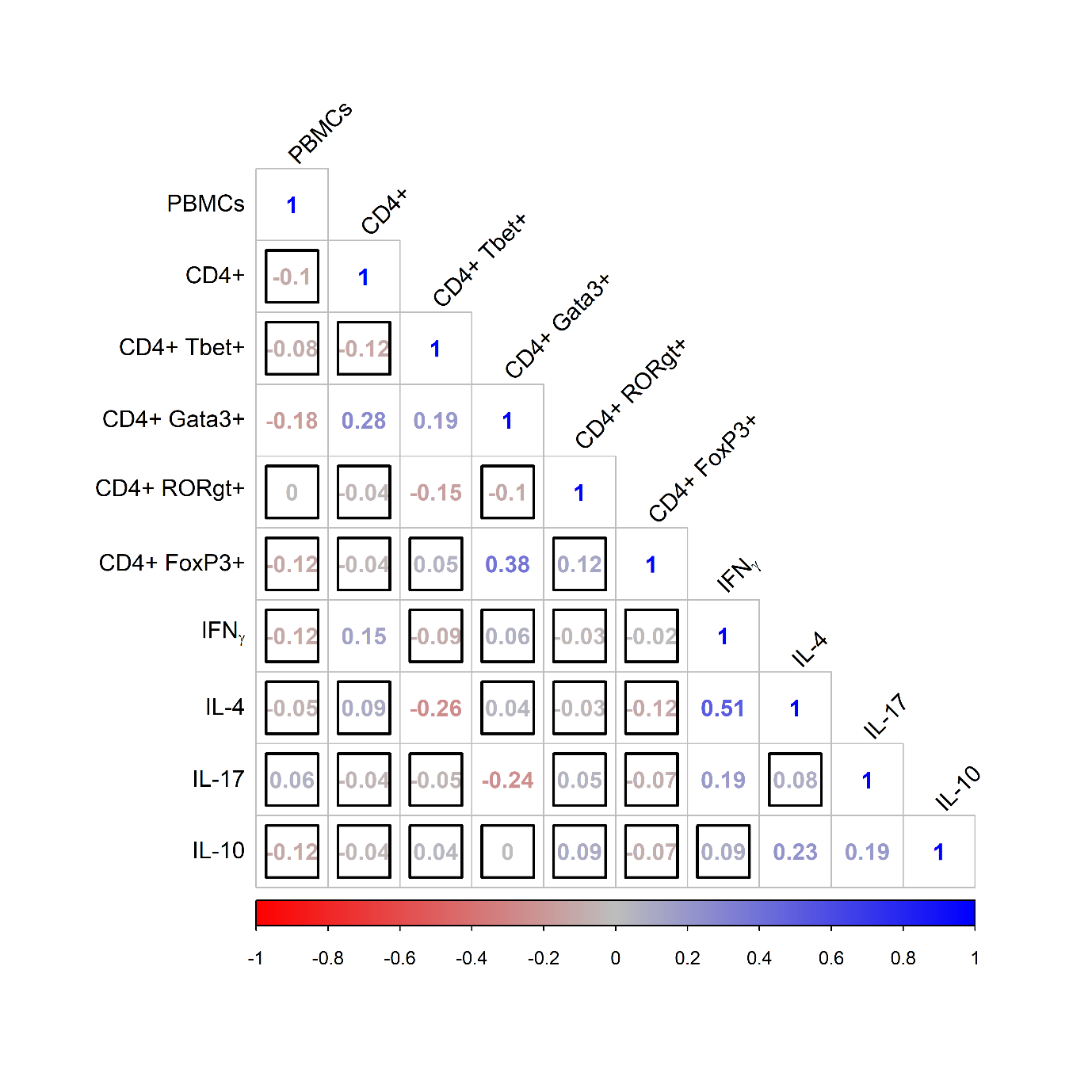

***Figure S1.*** *Correlation matrix showing Spearman’s rank correlations between pairs of immunological variables corrected for age and sex, where redder values indicate increasingly negative associations and bluer values indicate increasingly positive associations. Here, cell phenotypes are expressed as percentages of each cell type within the total leukocyte population rather than absolute numbers per ml blood; as such, compare this figure to Figure 3. Correlations outlined with a black square were not statistically significant at α=0.05.*

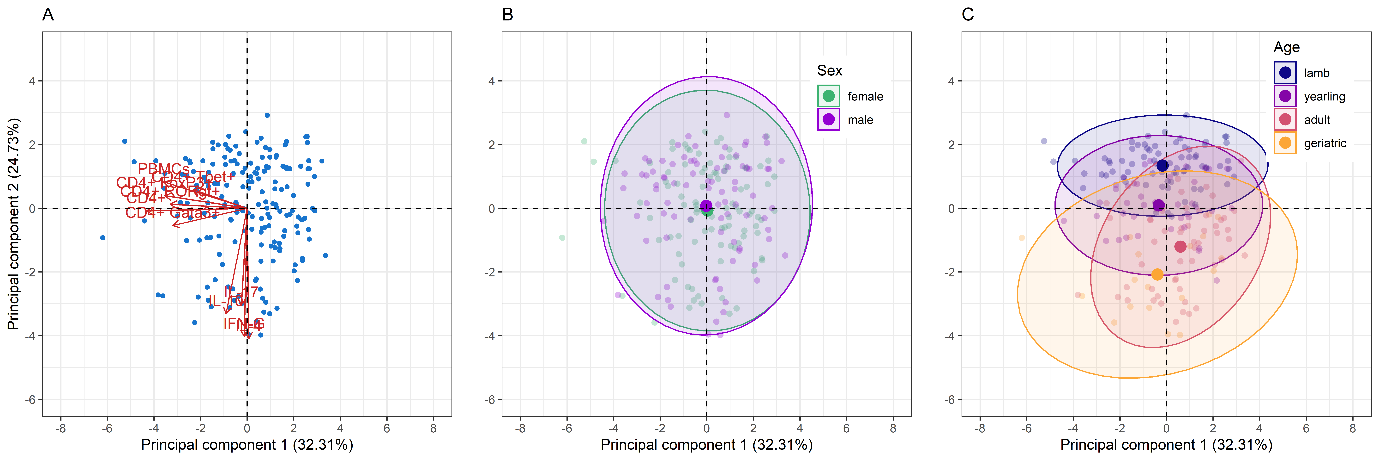

***Figure S2.*** *Biplots showing the results of principal components analysis (PCA) of 10 immunological variables. (A) Arrows show the loading of each variable on the first two principal components, with the length of the vector corresponding to the strength of the influence, while angles between vectors indicate the strength of association between the two variables. (B) and (C) show variation in the placing of data on the first two principal components by sex and age category respectively. Small points show each individual data point; large points show the centre of variation within each sex or age group; ellipses show 95% intervals around this.*
